# LRP1 mediates leptin transport by coupling with the short-form leptin receptor in the choroid plexus

**DOI:** 10.1101/2023.07.03.547520

**Authors:** Aaron Aykut Uner, Won-Mo Yang, Min-Cheol Kang, Kellen Cristina da Cruz Rodrigues, Ahmet Aydogan, Ji A Seo, Natalia F. Mendes, Min-Seon Kim, Fatima Ezzahra Timzoura, Michael J. Holtzman, Maria Lehtinen, Vincent Prevot, Young-Bum Kim

**Author notes:** These authors contributed equally (Aaron Aykut Uner, Won-Mo Yang, Min-Cheol Kang, Kellen Cristina da Cruz Rodrigues). Corresponding author: Young-Bum Kim, Ph.D. Division of Endocrinology, Diabetes, and Metabolism, Beth Israel Deaconess Medical Center 330 Brookline Avenue, Boston, MA 02215.

## Abstract

Adipocyte-derived leptin enters the brain to exert its anorexigenic action, yet its transport mechanism is poorly understood. Here we report that LRP1 (low-density lipoprotein receptor-related protein-1) mediates the transport of leptin across the blood-CSF barrier in *Foxj1* expressing cells highly enriched at the choroid plexus (ChP), coupled with the short-form leptin receptor, and LRP1 deletion from ependymocytes and ChP cells leads to leptin resistance and hyperphagia, causing obesity. Thus, LRP1 in epithelial cells is a principal regulator of leptin transport in the brain.

---

Leptin is an adipocyte-derived satiety factor that suppresses food intake and increases energy expenditure^1^. In order to achieve its physiological actions, it needs to enter the brain^2^. Until now, three potential gates for leptin entry into the brain are suggested as follows; (i) epithelial cells forming the blood-CSF barrier in the ChP^3^, (ii) endothelial cells forming the blood-brain barrier (BBB)^4^, and (iii) tanycytes in the mediobasal hypothalamus that form a barrier between the median eminence (ME) and the CSF through tight junctions^5^. The role of the long-form of leptin receptor (LepRb) in these routes for regulating body-weight homeostasis and energy balance is being uncovered ^6,7^. In particular, emerging evidence demonstrates that activation of the LepR in tanycytes of the ME is required for the regulation of leptin transport and energy balance^6^. Additionally, deleting LepR in brain endothelial and epithelial cells reduced leptin uptake by the brain and increased body weight under high-fat feeding^7^.

Interestingly, leptin also binds to the choroid plexus (ChP) in lean, leptin-deficient *ob/ob*, and leptin receptor (LepR)-deficient *db/db* mice as well as lean and obese Zucker rats with high affinity and high density^8^. These observations raise the possibility that the ChP may also be a major site for leptin transport through specific receptors for leptin other than the LepR^3,4,8^. In this respect, low-density lipoprotein receptor-related protein-1 or -2 (LRP1 or LRP2) has been suggested as a potential transporter for leptin^9,10^. However, the functional significance of LRP1 or LRP2 in the ChP for the regulation of body-weight homeostasis has not been elucidated.

We first determined the physiological role of LRP1 in the ChP in regulating body-weight homeostasis by studying mice lacking LRP1 in the ChP (Foxj1-Cre; LRP1*^loxP/loxP^*). The expression of Foxj1 is restricted to the epithelial cells of the ChP, where it is strongly expressed, and the ependymal cells of the ventricles, including the third ventricle (3V) (Supplementary Fig. 1a). LRP1 expression was abolished in the ChP of Foxj1-Cre; LRP1*^loxP/loxP^* mice, while its expression in other sites of the brain and peripheral organs was affected to a lower extent or not different between genotypes (Supplementary Fig. 1b). Interestingly, *in situ* hybridization for LRP1 and LepRa (the short-form of LepR) mRNA showed that the two transcripts were colocalized in the ChP (Supplementary Fig. 1c). At 18 weeks of age, the average body weights of male and female Foxj1-Cre; LRP1*^loxP/loxP^* mice were greatly increased by ∼20% and ∼35%, respectively, over LRP1*^loxP/loxP^* littermates (Fig. 1a and 1d). This effect was mainly attributed to an increased fat mass in these mice (Fig. 1b and 1e), confirmed by the direct dissection of fat depots (1c and 1f). Accordingly, serum leptin levels also markedly increased in Foxj1-Cre; LRP1*^loxP/loxP^*mice compared with LRP1*^loxP/loxP^* mice (Fig. 1h and 1m). These obesity phenotypes were most likely due to increased food intake (Fig. 1g and 1l). A rise in blood glucose and serum insulin levels was found in Foxj1-Cre; LRP1*^loxP/loxP^* mice (Fig. 1i–j and 1n–o). FFA levels didn’t change in these mice (Fig. 1k and 1p). Mice lacking LRP1 in the ChP were insulin resistant and glucose intolerant (Fig. 1r–u), evidenced by a significant decrease in AUCs (Supplementary Fig. 2a–d). In contrast, the deletion of LRP2 in the ChP had no effects on body weight, adiposity, and blood glucose (Supplementary Fig. 3c–e). This could be due to very little expression of LRP2 in epithelial cells of the ChP (Supplementary Fig. 3b). Together, these findings indicate that deficiency of LRP1 in the ChP may lead to an impairment of fuel metabolism and the accumulation of body fat.

**Fig. 1.**
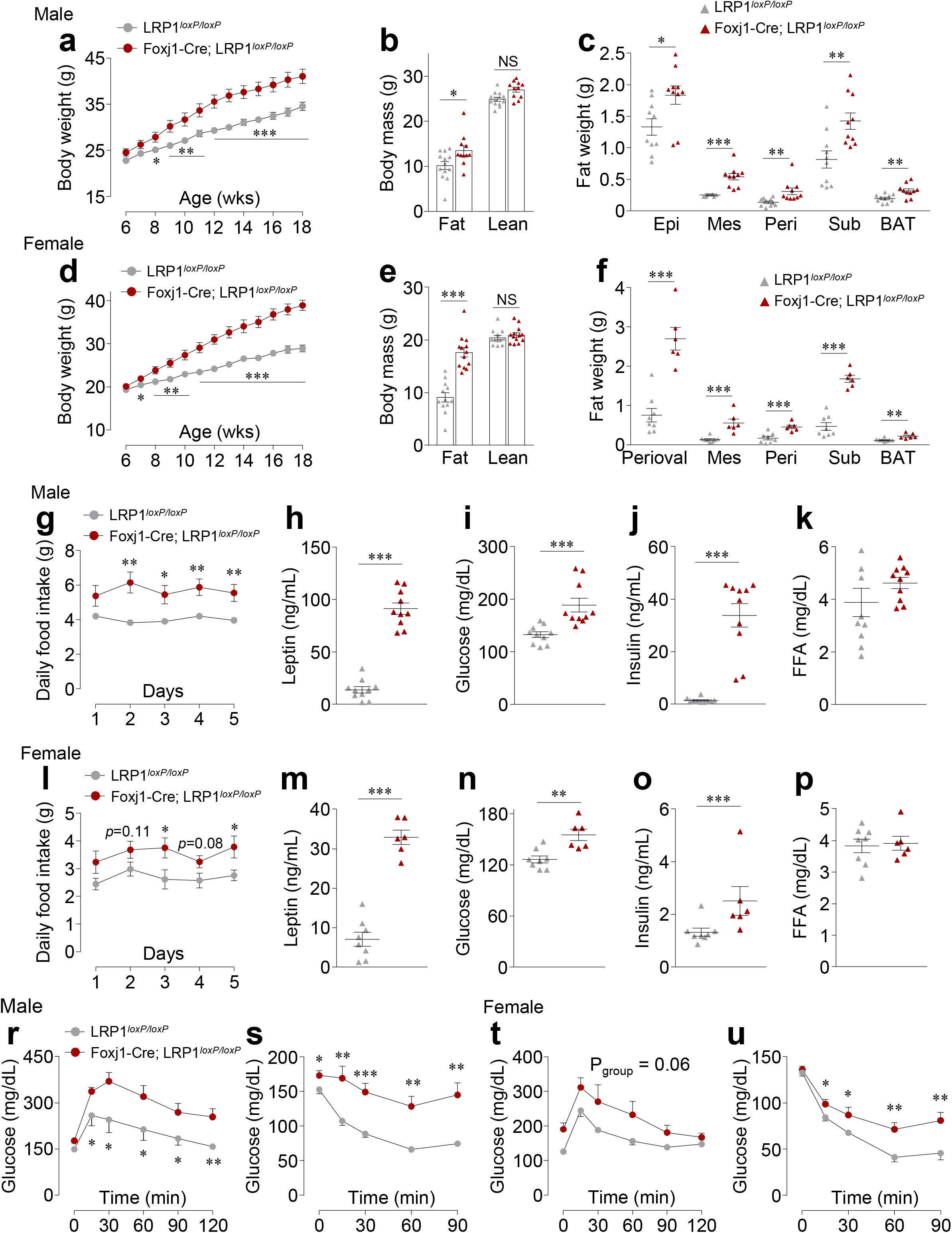
Loss of LRP 1 in the choroid plexus (ChP) leads to severe obesity phenotype. Body weight (n = 13 for LRP1*^loxP/loxP^*, n = 11 for Foxj1-Cre; LRP1*^loxP/loxP^*), (b) body mass (n = 13 for LRP1*^loxP/loxP^*, n = 11 for Foxj1-Cre; LRP1*^loxP/loxP^*), and (c) fat weight (n = 10 per group) were measured in male LRP1*^loxP/loxP^* and Foxj1-Cre; LRP1*^loxP/loxP^* mice. (d) Body weight (n = 12 for LRP1*^loxP/loxP^*, n = 13 for Foxj1-Cre; LRP1*^loxP/loxP^*), (e) body mass (n = 12 for LRP1*^loxP/loxP^*, n = 13 for Foxj1-Cre; LRP1*^loxP/loxP^*), and (f) fat weight (n = 9 per group) were measured in female LRP1*^loxP/loxP^*and Foxj1-Cre; LRP1*^loxP/loxP^* mice. (g) Daily food intake (n = 8 for LRP1*^loxP/loxP^*, n = 9 for Foxj1-Cre; LRP1*^loxP/loxP^*), (h) serum leptin (n = 8 for LRP1*^loxP/loxP^*, n = 6 for Foxj1-Cre; LRP1*^loxP/loxP^*), (i) blood glucose (n = 9 per group), (j) serum insulin (n = 8 for LRP1*^loxP/loxP^*, n = 6 for Foxj1-Cre; LRP1*^loxP/loxP^*), and (k) serum FFA (n = 8 for LRP1*^loxP/loxP^*, n = 6 for Foxj1-Cre; LRP1*^loxP/loxP^*) were measured in male LRP1*^loxP/loxP^* and Foxj1-Cre; LRP1*^loxP/loxP^*mice. (l) Daily food intake (n = 7 for LRP1*^loxP/loxP^*, n = 6 for Foxj1-Cre; LRP1*^loxP/loxP^*), (m) serum leptin (n = 10/group), (n) blood glucose (n = 10 per group), (o) serum insulin (n = 10 per group), and (p) serum FFA (n = 10 per group) were measured in female LRP1*^loxP/loxP^* and Foxj1-Cre; LRP1*^loxP/loxP^* mice. (r) Glucose tolerance test (GTT) (n = 6 for LRP1*^loxP/loxP^*, n = 8 for Foxj1-Cre; LRP1*^loxP/loxP^*) and (s) insulin tolerance test (ITT) (n = 7 for LRP1*^loxP/loxP^*, n = 9 for Foxj1-Cre; LRP1*^loxP/loxP^*) were performed in male LRP1*^loxP/loxP^*and Foxj1-Cre; LRP1*^loxP/loxP^* mice at 18–20 weeks of age. (t) Glucose tolerance test (GTT) (n = 6) and (u) insulin tolerance test (ITT) (n = 7 for LRP1*^loxP/loxP^*, n = 5 for Foxj1-Cre; LRP1*^loxP/loxP^*) were performed in female LRP1*^loxP/loxP^* and Foxj1-Cre; LRP1*^loxP/loxP^*mice at 30–32 weeks of age. Serum parameters, body mass, and fat weight were measured at 24 weeks of age for male mice and at 32 weeks of age for female mice. Daily food intake were measured at 18 weeks of age for male mice and at 32 weeks of age for female mice. Serum parameters were measured from overnight fasted mice. All bars and errors represent means ± SEM. *p* values by repeated measures two-way ANOVA in a, d, g, I, r–t, and u and by two-sided student *t*-test in b, c, e, f, h–k, and m–p are indicated. \**P* < 0.05, \*\**P* < 0.01, and \*\*\**P* < 0.001 vs. LRP1*^loxP/loxP^* mice.

To determine whether leptin enters the brain through LRP1 at the ChP to exert its anorexigenic action, peripheral or central leptin’s effect on food intake was assessed in mice lacking LRP1 in the ChP. As expected, food intake markedly decreased at 4–24 hours following the administration of intraperitoneal (IP) leptin in LRP1*^loxP/loxP^*mice (Fig. 2a). However, leptin’s anorexigenic effect was markedly impaired in all time points in Foxj1-Cre; LRP1*^loxP/loxP^* mice compared with control mice (Fig. 2a), suggesting that LRP1 in the ChP is required to regulate leptin transport. Importantly, in response to intracerebroventricular (ICV) leptin, the ability of leptin to suppress food intake was normal in Foxj1-Cre; LRP1*^loxP/loxP^*mice during 2–24 hours’ time points even though they are obese (LRP1*^loxP/loxP^*vs. Foxj1-Cre; LRP1*^loxP/loxP^*; 25.2 ± 0.53 (g) vs. 31.4 ± 0.85 (g), *P* < 0.0001) (Fig. 2b), indicating that LRP1 in the ChP mediates circulating leptin transport. Consistently, we also found that hypothalamic STAT3 phosphorylation significantly reduced in Foxj1-Cre; LRP1*^loxP/loxP^* mice or body weight-matched Foxj1-Cre; LRP1*^loxP/loxP^* mice compared with LRP1*^loxP/loxP^* mice when leptin was injected peripherally (Fig. 2c). This is further confirmed by immunoblotting analysis (Fig. 2e). In contrast, Foxj1-Cre; LRP1*^loxP/loxP^*mice displayed normal STAT3 phosphorylation of the hypothalamus by ICV leptin injection (Fig. 2d). Neither leptin-stimulated food intake nor hypothalamic STAT3 phosphorylation was altered in Foxj1-Cre; LRP2*^loxP/loxP^* mice (Supplementary Fig. 3f and 3g). These data are conflicting with the findings that LRP2 mediates leptin transport across the blood-CSF barrier^7^. Collectively, our data clearly demonstrate that LRP1 in the ChP could mediate leptin’s transport across the blood-CSF barrier at the ChP and thus play a pivotal role in regulating feeding behavior, underscoring a fundamental function for LRP1 in regulating leptin trafficking in the ChP. Moreover, these findings further imply that defective leptin trafficking in the ChP could contribute to the development of leptin resistance.

**Fig. 2.**
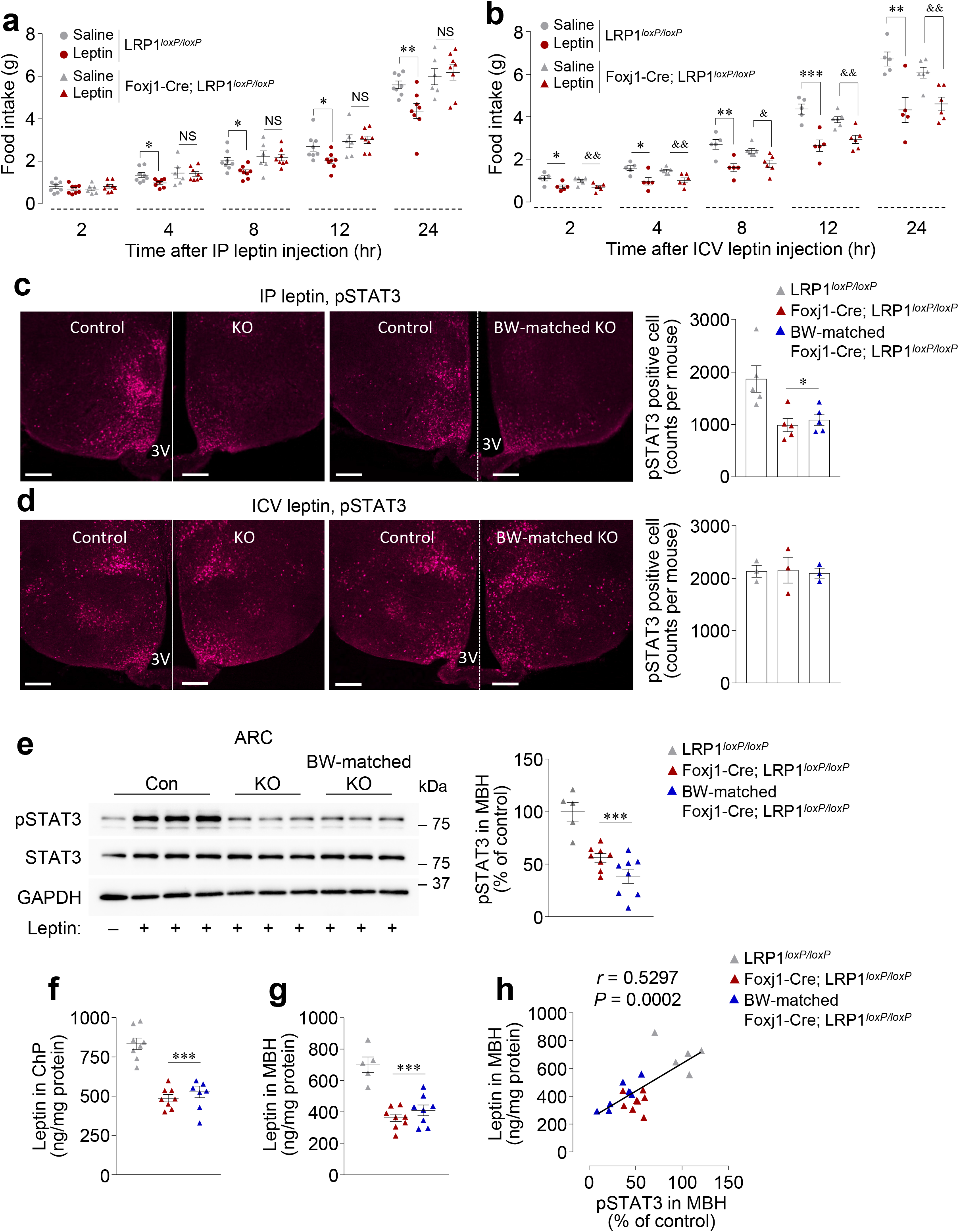
Control of leptin transport by LRP1 in the choroid plexus (ChP). Leptin-induced food intake was measured in LRP1*^loxP/loxP^* and Foxj1-Cre; LRP1*^loxP/loxP^* mice at 11–12 weeks of age (n = 8). Overnight-fasted mice were administered with IP leptin (3 mg/kg), and food intake was measured as indicated time points. (a) Leptin-induced food intake was measured in LRP1*^loxP/loxP^* and Foxj1-Cre; LRP1*^loxP/loxP^* mice at 14 weeks of age (n = 8). Overnight-fasted mice were administered with ICV leptin (1 μg/mouse), and food intake was measured as indicated time points. (b) STAT3 phosphorylation in the hypothalamus was measured in LRP1*^loxP/loxP^*, Foxj1-Cre; LRP1*^loxP/loxP^*, and body-weight (BM)-matched Foxj1-Cre; LRP1*^loxP/loxP^* mice at 18 weeks of age. Mice were administered with IP leptin (1 mg/kg) and sacrificed 30 min later. pSTAT3-positive neurons were detected by IHC analysis. Graph shows pSTAT3-positive cells from LRP1*^loxP/loxP^*, Foxj1-Cre; LRP1*^loxP/loxP^*, and body-weight (BM)-matched Foxj1-Cre; LRP1*^loxP/loxP^* mice (n = 5 per group). The scale bars represent 100 μm. (c) STAT3 phosphorylation in the hypothalamus was measured in LRP1*^loxP/loxP^*, Foxj1-Cre; LRP1*^loxP/loxP^*, and body-weight (BM)-matched Foxj1-Cre; LRP1*^loxP/loxP^* mice at 18 weeks of age. Mice were administered with intracerebroventricular (ICV) leptin (1 μg/mouse) and sacrificed 30 min later. pSTAT3-positive neurons were detected by IHC analysis. Graph shows pSTAT3-positive cells from LRP1*^loxP/loxP^*, Foxj1-Cre; LRP1*^loxP/loxP^*, and body-weight (BM)-matched Foxj1-Cre; LRP1*^loxP/loxP^* mice (n = 3 per group). The scale bars represent 100 μm. (d) STAT3 phosphorylation in the mediobasal hypothalamus (MBH) was measured in LRP1*^loxP/loxP^*, Foxj1-Cre; LRP1*^loxP/loxP^*, and body-weight (BM)-matched Foxj1-Cre; LRP1*^loxP/loxP^* mice at 18 weeks of age (n = 5–8 per group). Mice were injected with intraperitoneal (IP) leptin (1 mg/kg) and sacrificed 45 min later. MBH tissue lysates (20 µg) were separated by SDS– PAGE, and pSTAT3, STAT3, and GAPDH bands were visualized by immunoblotting. Bars show the densitometric quantitation of the pSTAT3 normalized to STAT3. (e) Leptin levels in the ChP were measured in LRP1*^loxP/loxP^*and Foxj1-Cre; LRP1*^loxP/loxP^*, and body-weight (BM)-matched Foxj1-Cre; LRP1*^loxP/loxP^* mice at 20 weeks of age (n = 5-8 per group). Mice were administered with intraperitoneal (IP) leptin (1 mg/kg) and sacrificed 30 min later. Leptin levels in ChP were determined by ELISA. (f) Leptin levels in the MBH were measured in LRP1^loxP/loxP^ and Foxj1-Cre; LRP1^loxP/loxP^, and body-weight (BM)-matched Foxj1-Cre; LRP1^loxP/loxP^ mice at 18 weeks of age (n = 5–8 per group). Mice were administered with intraperitoneal (IP) leptin (1 mg/kg) and sacrificed 45 min later. MBH tissues were lysed. Leptin levels in the MBH were determined by ELISA. (g) Relationship of leptin level with pSTAT3 in the MBH (n = 21) in LRP1^loxP/loxP^ and Foxj1-Cre; LRP1^loxP/loxP^, and body-weight (BM)-matched Foxj1-Cre; LRP1^loxP/loxP^ mice. *P* values were obtained by Spearman’s rank correlation analysis and *r* values indicate Spearman’s correlation coefficient. All experiments were performed with male mice. All bars and errors represent means ± SEM. *p* values by one-way ANOVA in a–f, and h are indicated. Post hoc analyses were done by the Bonferroni test for one-way and repeated measures two-way ANOVA. \**P* < 0.05, \*\**P* < 0.01, and \*\*\**P* < 0.001 vs. LRP1*^loxP/loxP^*mice or saline in the same group. ^&^*P* < 0.05 and ^&&^*P* < 0.01 vs. saline in the same group.

To investigate whether LRP1 is directly involved in leptin transport, we measured leptin levels in the ChP and mediobasal hypothalamus (MBH) after a bolus of IP leptin administration (1 mg/kg). Here we show that leptin levels in the ChP are significantly lower in Foxj1-Cre; LRP1*^loxP/loxP^* mice than LRP1*^loxP/loxP^* mice (Fig. 2f), indicating that less amount of leptin is traveled to the brain of Foxj1-Cre; LRP1*^loxP/loxP^* mice. This effect is unlikely due to body weight, as revealed by the fact that leptin levels in the ChP were comparable between Foxj1-Cre; LRP1*^loxP/loxP^* and BW-matched Foxj1-Cre; LRP1*^loxP/loxP^*mice (Fig. 2f). The immunoblotting analysis further confirmed these data (Supplementary Fig. 4). Similar results were also found in Foxj1-Cre; LRP1*^loxP/loxP^* mice when a sub-dose of leptin (0.5 mg/kg) was administered (Supplementary Fig. 4). Concurrently, we observed decreased leptin amounts in MBH in Foxj1-Cre; LRP1*^loxP/loxP^* mice after peripheral leptin stimulation (Fig. 2g), highlighting the necessity of LRP1 in the ChP in transporting circulating leptin. A positive correlation exists between leptin levels and pSTAT3 in the MBH during leptin stimulation (Fig. 2h), indicating that the degree of pSTAT3 in the MBH depends on the leptin amount delivered from the periphery. Our findings demonstrate that LRP1 in the ChP plays a role in delivering circulating leptin to the brain.

We explored the mechanisms underlying LRP1-mediated leptin transport. To address this, we tested the hypothesis that leptin binds to LRP1 and drives a physical interaction between LRP1 and the short-form of the leptin receptor LepRa in the ChP, and the heteromeric complex of an LRP1-LepRa undergoes endocytic internalization. The *in vivo* interaction of leptin or LepRa and LRP1 in response to leptin was measured using a proximity ligation assay (PLA)^11,12^. LRP1 deficiency in epithelial cells of the ChP didn’t lead to any abnormalities of cell morphology or apoptosis (Supplementary Fig. 5). Each far red spot represents a leptin-LRP1 interaction and a LepRa-LRP1 interaction (Fig. 3a). No far red spots were detected in a negative control that was performed the PLA with an IgG antibody (Supplementary Fig. 6). The PLA assay indicated that a leptin-LRP1 and LRP1-LepRa interaction in epithelial cells of the ChP in LRP1*^loxP/loxP^* mice notably increased in response to leptin stimulation, as evidenced by a marked elevation in far red spots (Fig. 3a). However, these effects were abolished when LRP1 was absent in the ChP (Fig. 3a). Quantification analysis revealed that leptin binding to LRP1 significantly decreased by ∼96% in Foxj1-Cre; LRP1*^loxP/loxP^* mice over LRP1*^loxP/loxP^*mice (Fig. 3b). In addition, leptin-induced interactions between LRP1 and LepRa markedly diminished by ∼74% in mice lacking LRP1 in the ChP (Fig. 3b). These data are further confirmed by *in vitro* study with ChP-derived epithelial Z310 cells showing that leptin interacts with LRP1 and increases the physical interaction of LRP1 and LepRa (Fig. 3c and 3d). Moreover, *in vitro* transient transfection studies showed strong binding of LRP1 to LepRa in cultured HEK293 and Neuro 2a cell lines (Fig. 3e and 3f).

**Fig. 3.**
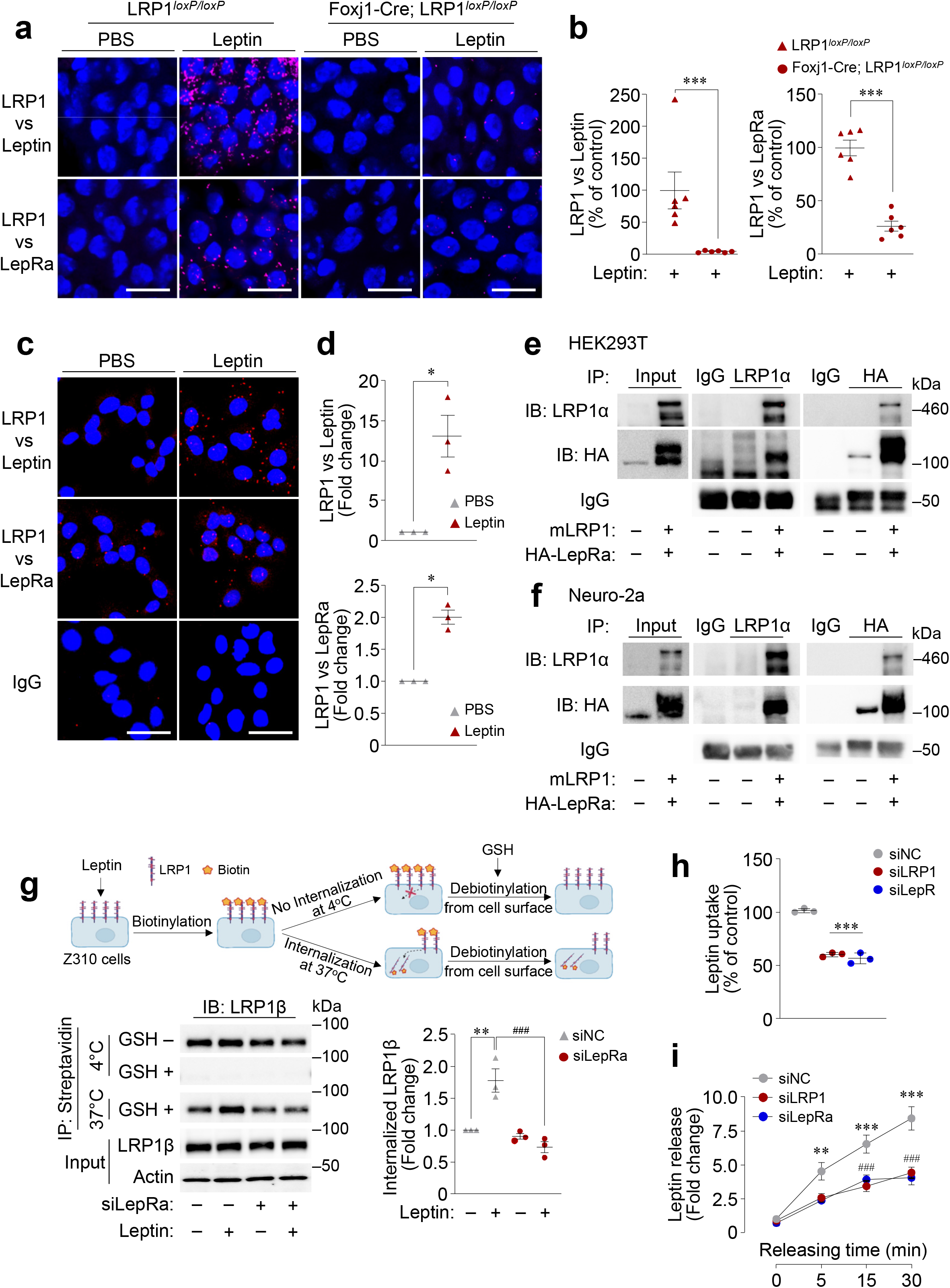
LRP1 mediates leptin transport via coupling with the LepR. (a) Exogenous leptin physically interacts with LRP1 and induces the physical interaction of LRP1 with LepRa in epithelial cells of the ChP in vivo. LRP1*^loxP/loxP^* and Foxj1-Cre; LRP1*^loxP/loxP^*mice at 18 weeks of age were administered with IP leptin (1 mg/kg) and sacrificed 30 min later. The choroid plexus (ChP) samples were harvested, and proximity ligation assays (PLA) was performed. Each far-red spot represents a leptin-LRP1 or a LRP1-LepR interaction. (b) Image data for leptin-LRP1 or LRP1-LepRa interactions were quantitated by Image J (n = 5–6). Nuclei were stained with DAPI (blue). The scale bar represents 20 μm. Data are represented as means ± SEM. \**P* < 0.05 and \*\**P* < 0.01 vs LRP1*^loxP/loxP^* mice by unpaired two-sided Student’s *t*-test. (c) Leptin induces the physical interaction of LRP1 and LepRa in Z310 cells. Z310 cells were stimulated with or without leptin (1 μM) for 10 min. PLA was performed. Each red spot represents a leptin-LRP1 or a LRP1-LepRa interaction. (d) Image data for leptin-LRP1 or LRP1-LepRa interactions were quantitated by Image J (n = 3). N represents an individual experiment. Nuclei were stained with DAPI (blue). The scale bar represents 25 μm. Data are represented as means ± SEM. \**P* < 0.05 and \*\**P* < 0.01 vs no leptin treatment by unpaired two-sided Student’s *t*-test. LRP1 interacts with LepR in (e) HEK 293 cells and (f) Neuro-2a cells. HEK 293 cells and Neuro-2a cells were transiently transfected with the plasmids of both pcDNA3.1(+)-C-DYK-mLRP1 (mLRP1) and pcDNA3.1(+)-N-HA-mLepRa (HA-mLepRa) for 48 h. The cell lysates (Input) were subjected to immunoprecipitation with either Normal rabbit IgG (IgG), LRP1α, or HA antibody, followed by immunoblotting with either LRP1α or HA antibody. The blots represent three independent experiments. (g) Leptin induces LRP1 internalization that is required for the LepRa. Z310 cells were transiently transfected with siNC or siRNA for LepRa (siLepRa). The cells were stimulated with or without leptin (0.5 μM) for 30 min, followed by cell surface biotinylation. The internalization of cell-surfaced receptors was induced by incubating the cells either at 4°C or 37°C, followed by debiotinylation with or without GSH buffer. The cell lysates were subjected to immunoprecipitation with streptavidin-conjugated beads. Biotinylated LRP1β (surface LRP1β and internalized LRP1β) and input LRP1β were visualized by immunoblotting (n = 3). Bars show densitometric quantitation of the internalized LRP1β (n = 3). N represents an individual experiment. Data are represented as means ± SEM. \*\**P* < 0.01 vs no leptin in siNC, ^###^*P* < 0.001 vs leptin in siNC by two-way ANOVA. (h) Leptin uptake in choroidal plexus epithelial Z310 cells. Rat choroidal plexus epithelial Z310 cells (Z310 cells) were transiently transfected with siRNA for negative control (siNC), siRNA for LRP1 (siLRP1), or siRNA for LepRa (siLepRa). The cells were stimulated with leptin (1 μM) for 15 min, and washed 5 times with ice-cold PBS. The cells were lysed with lysis buffer. The lysates were measured with leptin levels by ELISA (n = 3). N represents an individual experiment. Data are represented as means ± SEM. \*\*\**P* < 0.001 vs. siNC by one-way ANOVA. (i) Leptin release in choroidal plexus epithelial Z310 cells. Choroidal plexus epithelial cells take up and release leptin *in vitro*. Rat choroidal plexus epithelial Z310 cells (Z310 cells) were transiently transfected with siRNA for negative control (siNC), siRNA for LRP1 (siLRP1), or siRNA for LepRa (siLepRa). The cells were stimulated with leptin (1 μM) for 15 min, washed 5 times with ice-cold PBS and incubated with fresh leptin-free culture medium for 0, 5, 15, and 30 min. Culture medium was harvested as indicated time and leptin levels in culture medium (secreted leptin) were measured by ELISA (n = 3). N represents an individual experiment. Data are represented as means ± SEM. \*\**P* < 0.01 and \*\*\**P* < 0.001 vs. 0 min time-point in siNC, ^###^*P* < 0.001 vs 15 and 30 min time-point in siNC by two-way ANOVA.

We next assessed leptin-stimulated endocytic internalization of LRP1 by the cell surface protein biotinylation-based assay^13^. We observed that leptin rapidly stimulated LRP1 internalization by ∼1.8-fold over control but failed to induce LRP1 internalization in the absence of LepRa in ChP-derived epithelial Z310 (Fig. 3g). Accordingly, our findings further demonstrate that leptin uptake and release by ChP-derived Z310 epithelial cells were significantly diminished when LRP1 or LepRa was absent (Fig. 3h and 3i). These data suggest that LepRa is required for leptin-induced LRP1 internalization, an initial trigger step of leptin transport and further implicate that leptin transport is regulated by the mutual action of LRP1 and LepRa.

While our results implicating LRP1 in the ChP-mediated entry of leptin into the brain are quite intriguing, tanycytes, which have been convincingly demonstrated to transport leptin into the brain through a LepRb-mediated endocytotic mechanism^6^, are also enriched in Foxj1 (Figure Supplementary Fig. 8), thus contributing to the phenotype observed when LRP1 is deleted in these cells. Surprisingly, in ChP cells unlike tanycytes, it is LepRa, the short-form of LepR, but not the long-form of LepR^14^, which is widely expressed in epithelial cells and long-suspected to play a role in leptin transport^15^, that appears to associate with LRP1 upon stimulation with leptin. It is thus possible that leptin enters the brain at more than one route, through more than one mechanism or LepR isoforms, and with different target cell populations and physiological effects that need further studies to be teased apart. Finally, our genetic model involves the deletion of LRP1 in Foxj1-expressing cells even during development and not uniquely in adulthood, and it is thus not known how much of the metabolic phenotype might be due to neurodevelopmental alterations with long-term effects. Models in which LRP1 can be knocked out in more restricted cell populations and only during the experimental window help shed light on its role and the interaction of the multiple mechanisms of leptin transport that are in play.

In conclusion, we found that LRP1 in the ChP is an important regulator of leptin trafficking from circulation to the brain. Upon leptin stimulation, LRP1 physically interacts with LepRa in the plasma membrane of epithelial cells of the ChP, and the dimeric complexes of a LRP1-LepRa undergo co-endocytic internalization, which is an essential step for leptin transport. As a result, leptin enters the brain, ultimately leading to anorexigenic action. This model provides a new mechanism that advances our understanding of central leptin action and the pathogenesis of obesity.

## Methods

### Animal care

All animal care and experimental procedures were conducted in accordance with the National Institute of Health’s Guide for the Care and the Use of Laboratory Animals and approved by the Institutional Animal Care and Use Committee of Beth Israel Deaconess Medical Center. Unless otherwise stated, mice were allowed to access to standard chow (Teklad F6 Rodent Diet 8664, Harlan Teklad, Harlan, Madison, WI) and water provided *ad libitum* and they were housed at 22–24°C with a 12 h light-dark cycle with the light cycle starting from 6:00 a.m. to 6:00 p.m.

### Experimental animals

Mice bearing a *loxP*-flanked LRP1 allele (LRP1*^loxP/loxP^*) were purchased from The Jackson lab (Stock No: 012604, Bar Harbor, ME) and mice bearing a *loxP*-flanked LRP2 allele (LRP2*^loxP/loxP^*) were provided from Dr. Thomas Willnow (Max-Delbrueck-Center for Molecular Medicine, Berlin, Germany)^11^. Mice lacking LRP1 or LRP2 in Foxj1-expressing cells (Foxj1-Cre; LRP1*^loxP/loxP^*or Foxj1-Cre; LRP2*^loxP/loxP^*) were generated by mating LRP1*^loxP/loxP^* mice or LRP2*^loxP/loxP^* mice with Foxj1-Cre transgenic mice, respectively^16^. Foxj1-Cre; td-Tomato mice were generated by Foxj1-Cre mice mating with Cre-dependent td-Tomato reporter mice^17^ (gift from Dr. Brad Lowell). Genotypes of these mice were identified by polymerase chain reaction (PCR) or immunoblotting. All mice we studied are mixed background with 129 and C57BL/6.

### Body composition and food intake measurements

Mice were weighed from 5-6 weeks of birth and weekly thereafter. Fat and lean body mass were assessed using EchoMRI (Echo Medical Systems, Houston, TX, USA). Fat pads were harvested and weighed at the end of experiment in LRP1*^loxP/loxP^* (control) and Foxj1-Cre; LRP1*^loxP/loxP^* mice at 32 weeks of age. For the measurement of daily food intake, mice at 18 weeks of age were individually housed for 1 week prior to the measurement of food intake. A certain amount of food were given during both the adaptation period and experiment. Food intake was then measured over a 5-day period. Uneaten food was collected and measured. This amount was subtracted from the initial amount of food intake. Both male and female mice were used in this study.

### Blood parameter measurements

Blood was collected either from random-fed or overnight fasted mice via the tail. Blood glucose was measured using a OneTouch Ultra glucose meter (LifeScan, Inc., Milpitas, CA). Serum insulin and leptin levels were measured by ELISA (Crystal Chem, Elk Grove Village, IL). Serumfree fatty acid was measured by an enzymatic method (Wako Diagnostics, CA). The range and sensitivity for these parameters are glucose, 20–600 mg/dL; leptin, 1–25.6 ng/mL (sensitivity, 200 pg/mL using a 5 μL sample); insulin, 0.1–12.8 ng/mL (sensitivity, 0.1 ng/mL using a 5 μL sample); free fatty acid, 1–1000 mg/dL. All assays were performed blinded. Both male and female mice were used in this study.

### Glucose tolerance and insulin tolerance test

For the glucose tolerance test (GTT), male or female LRP1^loxP/loxP^ mice and Foxj1-Cre; LRP1^loxP/loxP^ mice at 20 weeks of age were fasted overnight, and blood glucose was measured before and 15, 30, 60, 90, and 120 min after an intraperitoneal injection of glucose (1.0 g/kg). For the insulin tolerance test (ITT), male or female LRP1^loxP/loxP^ mice and Foxj1-Cre; LRP1^loxP/loxP^ mice at 18 weeks of age were fasted for 5 h and blood glucose was measured before and 15, 30, 60, and 90 min after an intraperitoneal injection of human insulin (0.75 IU/kg of body weight; Humulin R, Eli Lilly, Indianapolis, IN). The area under the curve for GTT and ITT was calculated^18^.

### Leptin levels in the choroid plexus (ChP) and serum

Male LRP1^loxP/loxP^ and Foxj1-Cre; LRP1^loxP/loxP^ mice were injected with IP leptin (1 mg/kg) or saline. Thirty min after the injections, the ChP and serum were rapidly harvested. Leptin levels were measured by ELISA or immunoblotting.

### Leptin-induced food intake

For the measurement of intraperitoneal (IP) leptin-induced food intake, male LRP1^loxP/loxP^ and Foxj1-Cre; LRP1^loxP/loxP^ mice at 14 weeks of age were individually housed for at least 1 week. Mice were fasted overnight and administered with IP leptin (3 mg/kg, National Hormone and Peptide Program, Torrance, CA) or saline. For the measurement of intracerebroventricular (ICV) leptin-induced food intake, a 26-gauge stainless steel cannula (Protech International, TX) was implanted into the lateral ventricle of male mice LRP1*^loxP/loxP^* and Foxj1-Cre; LRP1*^loxP/loxP^* mice at 14 weeks of age (the coordinates from the Bregma: − 0.46 mm antero-posterior, + 1 mm medio-lateral and - 2.2 dorso-ventral) under deep anesthesia. A Bregma-Lambda correction (up to 0.07 mm) was done before the cannula implantation. Mice were individually housed for at least 1 week for recovery. After the recovery period, mice were handled daily for 3 days to acclimatize them to the injection procedure. Mice were then fasted overnight and administered with ICV leptin (1 μg/mouse) or saline. Food intake was measured 2, 4, 8, 12, and 24 h after the injections of IP or ICV leptin or saline.

### Leptin-Induced STAT3 phosphorylation

Male LRP1^loxP/loxP^ and Foxj1-Cre; LRP1^loxP/loxP^ mice were injected with IP (1 mg/kg) or ICV leptin (1 μg/mouse) and transcardially perfused with saline 30 min after the leptin injection. Brains were extracted and incubated in 10% formalin overnight for extended fixation. The brains were then incubated in 20% sucrose and sectioned coronally on a sliding microtome (Leica SM 2000R, IL) at 35 µm (4 series equally). After washing in PBS (3 times, 5-min each), the serial brain sections were incubated in 1% NaOH + 1% H_2_O_2_ (20 min), 0.3% glycine (10 min) and 0.03% SDS (10 min). The sections were blocked in 10% normal donkey serum for 1 h at room temperature (RT) and then incubated overnight at RT in the same blocking solution containing pSTAT3 antibody (Phospho-STAT3 (Tyr705), 1:500, Cell Signaling Technology, Danvers, MA). The next day, the sections were washed in PBS (3 times, 5-min each) and then incubated in the same blocking solution containing secondary antibody (Donkey anti-rabbit Alexa fluorophore 647, 1:500 Thermo Fisher Scientific, Waltham, MA) for 1 hour at RT. After the incubation, the sections were washed in PBS (3 times, 5-min each) and mounted onto slides. The images were taken with a slide scanner microscope (Olympus VS120, Tokyo, Japan)^19^. The pSTAT3 positive cells in the hypothalamus from 3-4 sections per mouse were quantified with Image J program (ImageJ, U. S. National Institutes of Health, MD).

### Histopathology

Mice were transcardially perfused with PBS followed by a 10% formalin solution. The brain samples were further fixed in a 10% neutral formalin solution overnight and processed using graded alcohols and xylene. For histopathological examination, sections (5 μm) were taken from paraffin-embedded brain tissue samples, stained with hematoxylin-eosin (H&E) (MHS32, Sigma-Aldrich, Louis, MO) and examined under a light microscope (Olympus BH-2, Shinjuku, Tokyo, Japan).

### Immunohistochemistry (IHC)

IHC analysis was performed on 5 μm paraffin sections. Sections were stained with Caspase-3 antibody (dilution; 1:500, cat#: ab184787, Abcam, Waltham, MA) using the standard avidin– biotin– peroxidase complex (ABC) method according to the manufacturer’s instructions (Mouse and Rabbit Specific HRP/DAB (ABC) Detection IHC kit, Cat#: ab64264, Abcam). They were dewaxed in xylene and subsequently passed through in ethanol (100%, 80% and 50%). Antigen retrieval was applied by microwaving the sections in citrate buffer (pH = 6.0) at a sub-boiling temperature for 10 min. After a second protein blocking, sections were incubated with primary antibody overnight at 4°C. The brown color of immunopositivity in sections was improved with 3,3’-diaminobenzidine tetrahydrochloride (DAB-H_2_O_2_) substrate in PBS for 5 min. Slides were counterstained with Mayer’s hematoxylin (Sigma-Aldrich), dehydrated and mounted with Entellan (Sigma-Aldrich).

### TUNEL

Terminal deoxynucleotidyl transferase-mediated dUTP nick end-labeling (TUNEL) assay was performed on paraffin embedded tissue sections according to the manufacturer’s instructions to detect DNA fragmentation in apoptosis (TUNEL assay kit-HRP-DAB, Cat#: ab206386, Abcam). Briefly, the sections were dewaxed, hydrated and subsequently were digested with Proteinase K (20 mg/ml) for 20 min at RT. Endogenous peroxidase activity was quenched with 3% H_2_O_2_ for 5 min. The slides were incubated with terminal deoxynucleotidyl transferase (TdT) Labeling Reaction Mix in a humidified chamber at 37°C for 1.5 h. The sections were covered with stop buffer solution at room temperature for 5 min to terminate the labeling reaction. The reaction color was developed by DAB and the sections were counterstained with methyl green. The slides were then examined under a light microscope (Nikon Eclipse E600, Nikon, Melville, NY).

### Cell culture

Rat choroid plexus-derived epithelial Z310 cells (a gift from Dr. Wei Zheng, Purdue University, West Lafayette, IN) were cultured with DMEM/F-12 supplemented with 10% FBS, 1% antibiotic-antimycotic, gentamicin (40 μg/ml), and EGF (10 ng/ml). Neuro-2a mouse neuroblasts (American Type Culture Collection, Manassas, VA) were cultured with EMEM (American Type Culture Collection) with 10% FBS, and 1% antibiotic-antimycotic. HEK293T human embryonic kidney cells were cultured with high glucose DMEM supplemented with 10 % FBS and 1% antibiotic-antimycotic. All reagents for cell culture were purchased from Thermo Fisher Scientific (Waltham, MA).

### Transfection

Z310 cells were transiently transfected with small interfering RNA (siRNA) using Lipofectamine RNAiMax (Thermo Fisher Scientific) according to the manufacturer’s instructions. Cells were used for experiments 48 h after transfection. The *LRP1* siRNA (Cat#: s152489), *LepR* siRNA (Cat#: s217853), and negative control siRNA (Cat#: 4390844) were purchased from Thermo Fisher Scientific. Neuro-2a cells or HEK293 cells were transiently transfected with pcDNA3.1(+)-C-DYK-mLRP1 (mouse LRP1 CDS; NM_008512.2, Genscript) and pcDNA3.1(+)-N-HA-mLepR (mouse LepR isoform A CDS; NM_001122899.2, Genscript) using Lipofectamine 3000 (Thermo Fisher Scientific) according to the manufacturer’s instructions. Cells were used for experiments 48 hours after transfection.

### Leptin release assay

For leptin release assay, Z310 cells were treated with leptin (1 μM, R&D Systems, Minneapolis, MN) for 15 min, washed three times with cold PBS and incubated with 37°C pre-heated medium for 5, 15, and 30 min. The medium was harvested at each time points and immediately frozen in dry ice. Leptin concentrations in the medium were measured by ELISA (Crystal Chem).

### LRP1 internalization assay

Z310 cells were treated with leptin (0.5 μM) for 30 min at 4°C and washed three times with cold PBS. Cells were then biotinylated with EZ-Link™ Sulfo-NHS-SS-Biotin (0.3 mg/ml, Thermo Fisher Scientific) for 30 min at 4°C, and washed two times with cold PBS containing glycine (20 mM) for 5 min. To stimulate endocytosis, cells were incubated for 2 min at 37°C. For the removal of unendocytotic biotin from the cell surface, glutathione (GSH) buffer (50 mM reduced GSH, 75 mM NaCl, 75 mM NaOH, 5 mM EDTA) was incubated 2 times for 15 min at 4°C and quenched by adding iodoacetamide (50 μg/ml) for 10 min. Cells were lysed with NP-40 lysis buffer (Thermo Fisher Scientific) supplemented with 1X Halt™ Protease and Phosphatase Inhibitor Cocktail (Thermo Fisher Scientific) and PMSF (1 mM, Merck). Cell lysates (20 μg) were subjected to immunoprecipitation with Pierce™ Streptavidin Magnetic Beads (10 μl, Thermo Fisher Scientific) overnight at 4°C, followed by immunoblotting. LRP1β bands were visualized with SuperSignal Chemiluminescent Substrates (Thermo Fisher Scientific) and quantitated by Image J program (NIH, Bethesda, MD).

### Immunoprecipitation analysis

For interaction of LRP1 and LepR, cell lysates (500 μg) were subjected to immunoprecipitation with LRP1α (1 μg, Cat#: L2295, Merck, Rahway, NJ), HA (0.2 μg, Cat#: 3724, Cell Signaling Technology, Danvers, MA) or normal rabbit IgG (Cat#: 2729, Cell Signaling Technology) coupled to Dynabeads™ Protein A (20 μl, Thermo Fisher Scientific) overnight at 4°C. The immunoprecipitants were washed four times with lysis buffer, and then were eluted by 1X Novex™ Tris-Glycine SDS sample buffer (Thermo Fisher Scientific) with 1X NuPAGE™ Sample Reducing Agent (Thermo Fisher Scientific) for 10 min at 100°C.

### Immunoblotting analysis

Cell or tissue lysates (10–30 μg protein) or immunoprecipitated samples were resolved by SDS–PAGE and transferred to nitrocellulose membranes (Bio-Rad Laboratories, Hercules, CA). The membranes were incubated with antibodies against leptin (Cat#: 500-P68, PeproTech, rebury, NJ), pSTAT3 (Cat#: 9145, Cell Signaling Technology), STAT3 (Cat#: 9132, Cell Signaling Technology), LRP1α (Cat#: L2295, Merck), LRP1β (Cat#: ab-92544, Abcam, Waltham, MA), HA (Cat#: 3724, Cell Signaling Technology), GAPDH (Cat#: sc-47724, Santa Cruz Biotechnology, Dallas, TX), or β-Actin (Cat#: A2228, Merck). The membranes were washed and incubated with secondary antibodies (Cat#: 7074 or 7076, Cell Signaling Technology). The bands were visualized with SuperSignal Chemiluminescent Substrates (Thermo Fisher Scientific) and quantitated by Image J program (NIH, Bethesda, MD).

### *In situ* proximity ligation assay (PLA)

The choroid plexus (ChP) was harvested from the lateral ventricle of overnight-fasted male LRP1^loxP/loxP^ and Foxj1-Cre; LRP1^loxP/loxP^ mice 30 min after IP leptin injection (1 mg/kg). The ChP samples were placed onto slides and fixed with ice-cold methanol for 15 min at −20°C. Antigen retrieval was achieved by incubating the samples in citrate buffer (pH: 6.0) for 30 min at 60°C followed by Duolink^®^ In Situ PLA^®^ blocking solution for 1 h at 37°C and incubated with primary antibodies against LRP1(Cat#: sc-57353, Santa Cruz Biotechnology), leptin (Cat#: PA5-47023, Thermo Fisher Scientific) and LepR (Cat#: PA1-053, Thermo Fisher Scientific) overnight at 4°C. PLA was performed using Duolink^®^ In Situ PLA^®^ Probes and Duolink^®^ In Situ Detection Reagents FarRed (Sigma-Aldrich). The nuclei of ChP samples were stained using Duolink^®^ In Situ Mounting Medium with DAPI (Sigma-Aldrich). For *in vitro* studies, Z310 cells were incubated with or without leptin (100 nM) for 10 min. Cells were washed with ice-cold PBS three times and fixed with ice-cold methanol for 10 min at −20°C. Cells were incubated with primary antibodies against leptin (Cat#: PA5-47023, Thermo Fisher Scientific), LRP1 (Cat#: 37-3800, Thermo Fisher Scientific), or leptin receptor (Cat#: PA1-053, Thermo Fisher Scientific) overnight at 4°C. PLA was performed using the NaveniFlex GM or NaveniFlex RM kit (Navinci, Uppsala, Sweden). The nuclei of cells were stained using Duolink^®^ In Situ Mounting Medium with DAPI. Images from both *in vivo* and *in vitro* studies were captured by the Upright Confocal System (ZEISS LSM 880, ZEISS, Dublin, CA) and image data were analyzed by ZEISS ZEN lite software (ZEISS) or Image J program (NIH).

### Quantitative reverse transcription-polymerase chain reaction (qRT-PCR)

Total RNA was extracted using a TRIzol reagent (Thermo Fisher Scientific) and synthesized cDNA with a High-Capacity cDNA Reverse Transcription Kit (Thermo Fisher Scientific). qRT-PCR was performed with 20 ng of cDNA, the LepR or β-Actin primers, and a Power SYBR Green PCR Master Mix (Thermo Fisher Scientific) using a QuantStudio 6 (Thermo Fisher Scientific). Relative LepR mRNA expression levels were determined using the 2^−ΔΔCT^ method normalized to β-Actin mRNA expression levels. For LepR mRNA, the forward primer 5’-CTTTTCTGTGGGCAGAATCAGC-3’ and the reverse primer 5’-AGCACTGAGTGACTGCACAGCA-3’ were used. For β-Actin mRNA, the forward primer 5’-CATTGCTGACAGGATGCAGAAGG-3’ and the reverse primer 5’-TGCTGGAAGGTGGACAGTGAGG-3’ were used.

### RNAscope fluorescence *in situ* hybridization

Fluorescence in situ hybridization (FISH) was performed on fresh-frozen brain sections of the ChP (18µm) of adult male mice using the RNAscope Multiplex Fluorescent Kit v.2, according to the manufacturer’s instructions (Advanced Cell Diagnostics). Specific probes were used to detect LepR variant 3 (496901-C3, NM_001122899.1, target region 3291–4713, i.e., LepRa) and LRP1 (465231, NM_008512.2, target region 495 - 1451) mRNAs.

### Statistical analysis

The data were checked for normality and homogeneity of variance by Shapiro-Wilk and Levene’s tests, respectively. Logarithmic transformation was done for the data that violate the assumptions of normality and homogeneity of variance. Student’s *t*-tests were used throughout the study to compare two distinct groups. One-way analysis of variance (ANOVA) with the Bonferroni post hoc test was done to compare three groups. Two-way ANOVA was conducted for leptin uptake and LRP1 internalization experiments. Repeated measures two-way ANOVA was performed for body weight, daily food intake, GTT, ITT, and leptin-induced food intake experiments. When the main effect or interaction was significant by two-way ANOVA, post hoc analyses were further performed by general linear model procedures in SPSS or the Bonferroni test in Prizm. Statistical significances were analyzed using Prizm 7.0 software (GraphPad Software, La Jolla, CA) and SPSS version 22.0 for window (SPSS, Inc., Chicago, IL). Differences were considered significant at *P* ≤ 0.05. Data are presented as the mean or individual values ± SEM.

## ACKNOWLEDGEMENTS

This work was supported by grants from the National Institutes of Health (Health (R01DK111529, R01DK106076, and R01DK123002 to Y.B.K.) and the Basic Science Research Program through the National Research Foundation of Korea (NRF) funded by the Ministry of Education (NRF-2022R1A2C1013479 to J.A.S.). K.C.R. is a recipient of São Paulo Research Foundation from Brazil (FAPESP 2019/19938-5). We thank Brad Lowell for providing tdTomato mice; Thomas Willnow for providing LRP2 floxed mice; Wei Zheng for providing rat choroid plexus cell line Z310; Navinci for providing the PLA kits, Thiago Araujo for technical help and Brad Lowell, Barbara Kahn, Mark Anderson, all members of the Kim lab for their helpful advice and discussion. We are grateful to the Histology Core at Beth Israel Deaconess Medical Center.

## AUTHOR CONTRIBUTIONS

A.A.U., W.M.Y., M.C.K., K.C.R., A.A., J.A.S., N.F.M., M.S.K., M.L., V.P., and Y.B.K. designed and performed experiments. A.A.U. and W.M.Y. conducted interaction and binding assays. F.E.T. performed the RNAscope experiments. M.J.H. provided key material and expertise. All authors analyzed and interpreted experimental data. A.A.U., W.M.Y., V.P., and Y.B.K. wrote the manuscript with input all other authors.

## Supplementary Figures

**Supplemental Fig. 1.**
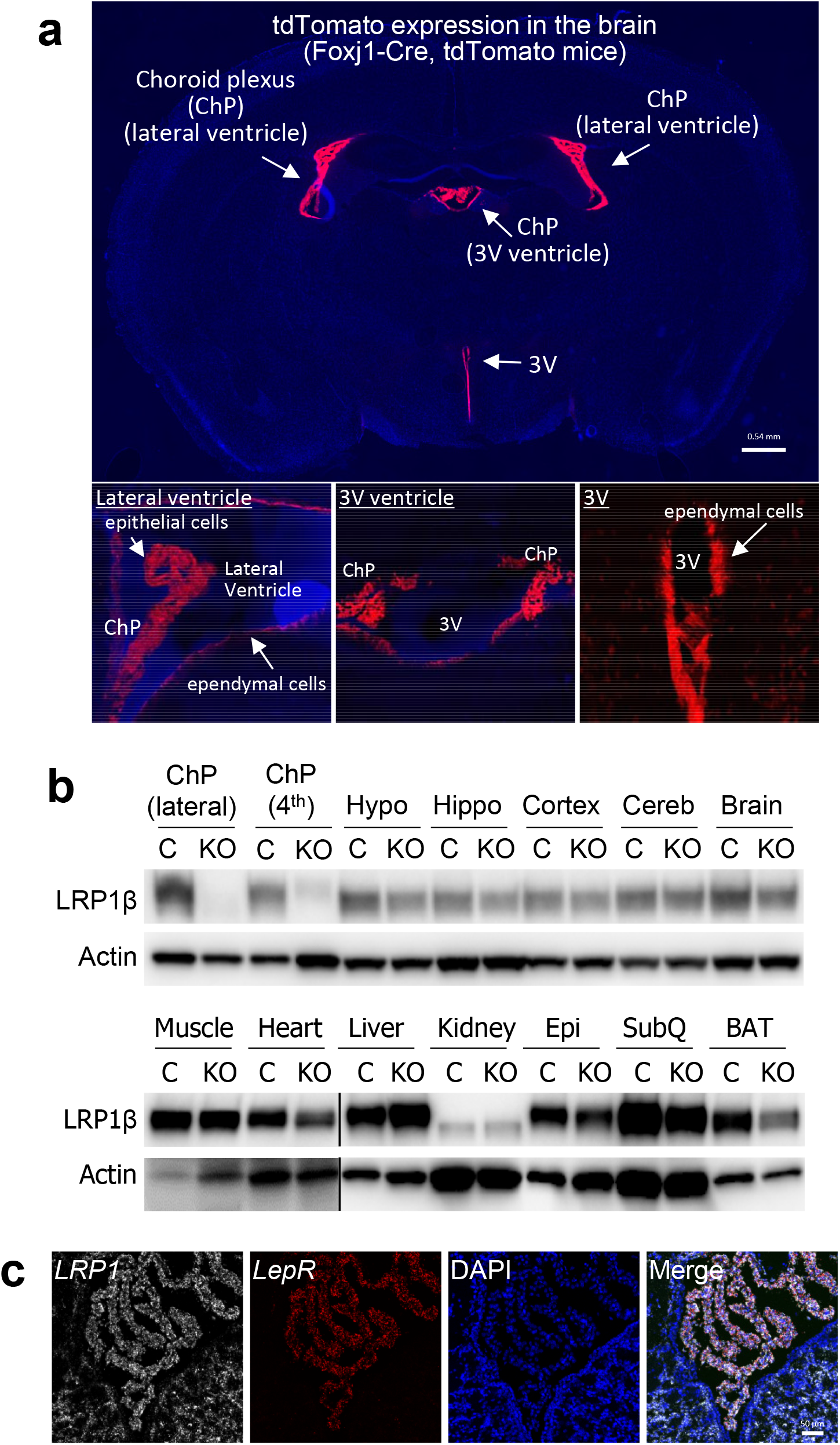
LRP1 expression in the choroid plexus (ChP) and multiple organs. (a) TdTomato expression in the brain of Foxj1-Cre, td-Tomato mice. The expression of tdTomato is restricted to the epithelial cells of the ChP and the ependymal cells of the ventricles. (b) LRP1 expression in multiple tissues of Foxj1-Cre; LRP1*_loxP/loxP_* mice. Tissue lysates (10–50 µg) were separated by SDS–PAGE, and LRP1 and actin bands were visualized by immunoblotting. ChP (lateral): choroid plexus lateral ventricle, ChP (4_th_): choroid plexus 4_th_ ventricle Hypo: hypothalamus, Hippo: hippocampus, Cereb: cerebellum, Epi: epididymal, BAT: brown adipose tissue, SubQ: subcutaneous fat. Lines in the blots indicates different blots (c) LRP1 and LepR expression in the ChP. RNA scope was performed. White indicates LRP1 and red indicates LepR.

**Supplemental Fig. 2.**
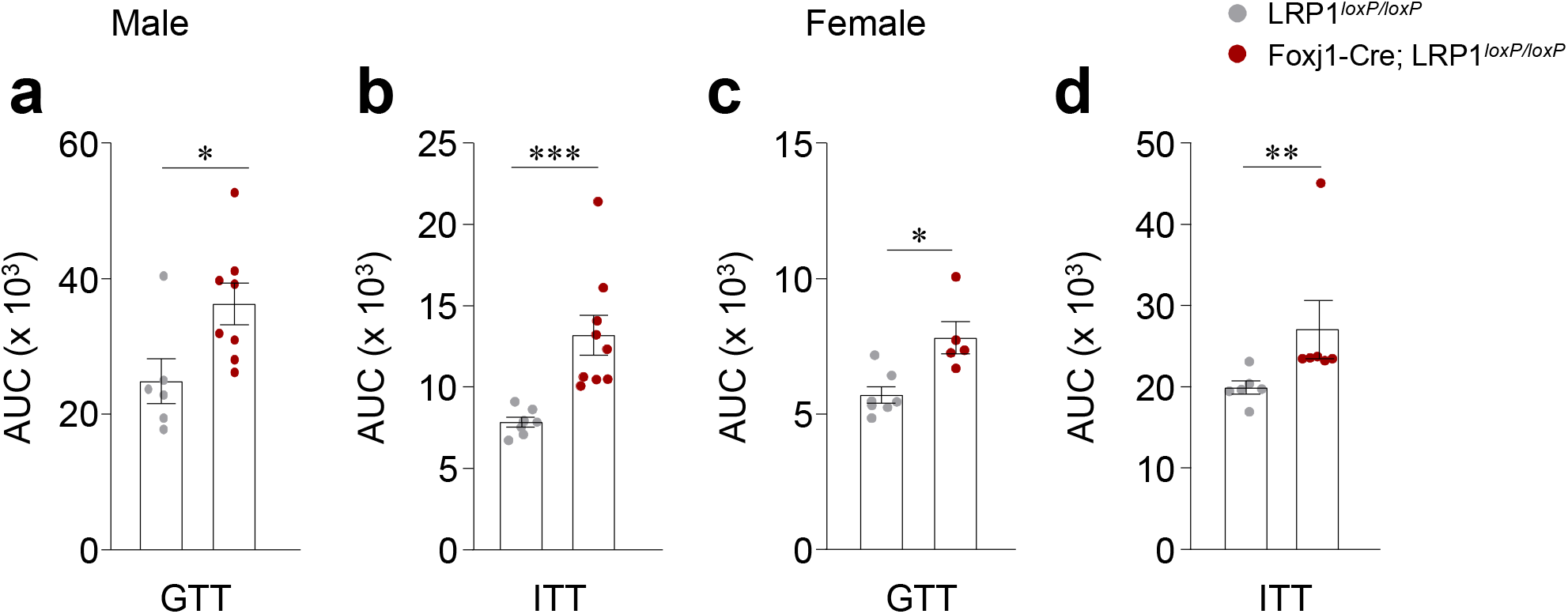
Area under the curves (AUCs) from GTT and ITT experiments in Fig. 2r–u (n = 5–9 mice per group) were calculated. All bars and errors represent means ± SEM. \**P* < 0.05, \*\**P* < 0.01, and \*\*\**P* < 0.001 vs. LRP1*_loxP/loxP_* mice by one-sided student *t*-test.

**Supplemental Fig. 3.**
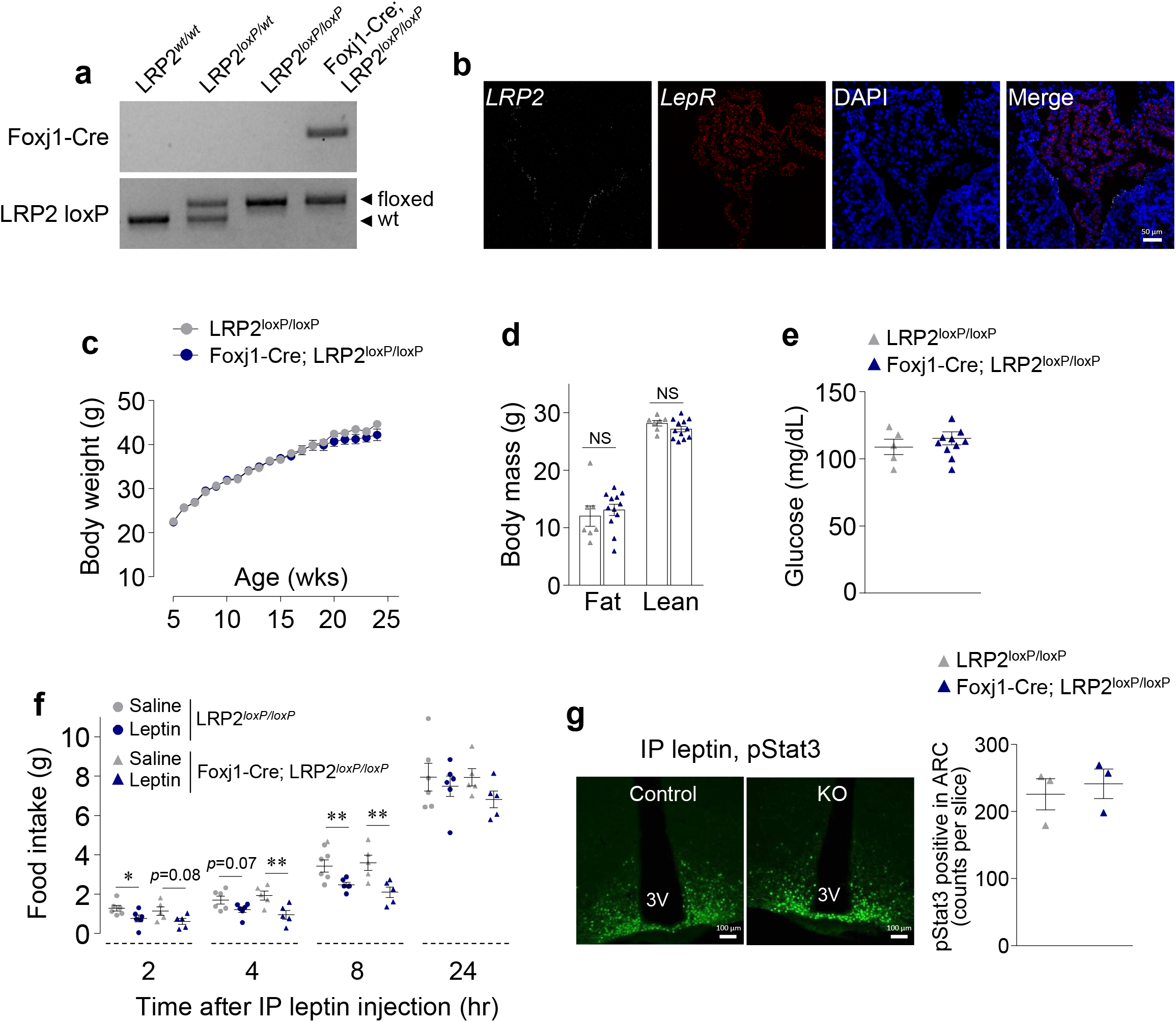
Deletion of LRP2 from the choroid plexus (ChP) does not affect body weight and leptin-induced food intake. (a) Genotyping, (b) RNAscope, (c) body weight, (d) body mass, (e) blood glucose, (f) leptin-induced food intake, and (g) pSTAT3 in LRP2*_loxP/loxP_* and Foxj1-Cre; LRP2*_loxP/loxP_* male mice. Body mass was measured by an MRI at 20 weeks of age. Glucose levels were measured from overnight fasted mice at 26 weeks of age. Leptin-induced food intake was measured at 24 weeks of age. pStat3-positive neurons were detected by IHC analysis. All bars and errors represent means ± SEM. \**P* < 0.05 and \*\**P* < 0.01 vs. LRP1*_loxP/loxP_* mice by repeated measures two-way ANOVA.

**Supplemental Fig. 4.**
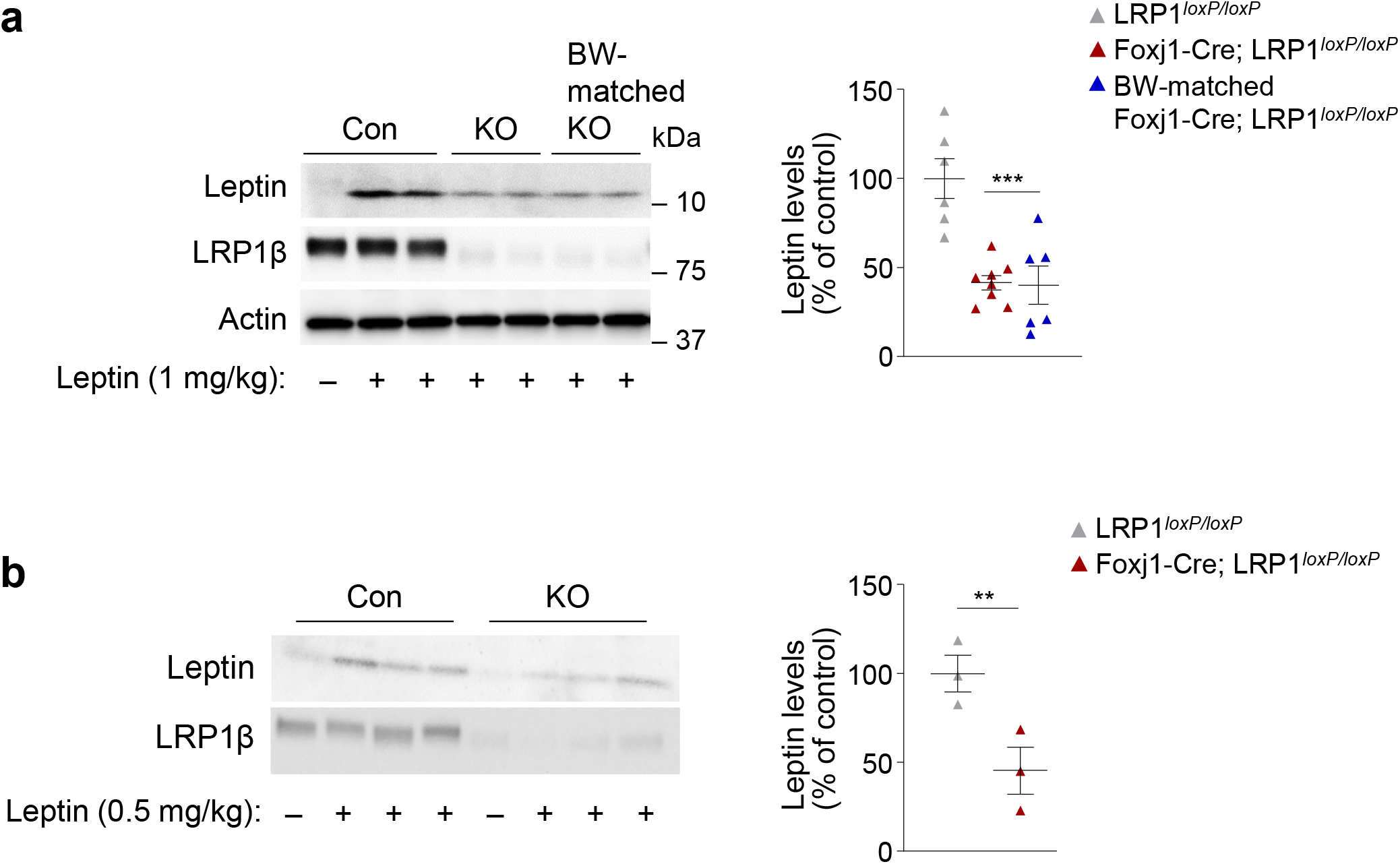
(a) Leptin levels in the choroid plexus (ChP) were measured in LRP1*_loxP/loxP_* and Foxj1-Cre; LRP1*_loxP/loxP_*, and body-weight (BM)-matched Foxj1-Cre; LRP1*_loxP/loxP_* mice at 20 weeks of age. Mice were administered intraperitoneal (IP) leptin (1 mg/kg) and sacrificed 30 minutes later. Tissue lysates (10 µg) were separated by SDS–PAGE. Leptin, LRP1β, and Actin bands were visualized by immunoblotting. The graph shows densitometric quantitation of immunoblot leptin from LRP1*_loxP/loxP_*, Foxj1-Cre; LRP1*_loxP/loxP_*, and body-weight (BM)-matched Foxj1-Cre; LRP1*_loxP/loxP_* mice (n = 6–8 per group). All bars and errors represent means ± SEM. *P* values by one-way ANOVA are indicated. \*\*\**P* < 0.001 vs. LRP1*_loxP/loxP_* mice in the same group. (b) Leptin levels in the choroid plexus (ChP) were measured in LRP1*_loxP/loxP_* and Foxj1-Cre; LRP1*_loxP/loxP_*, and body-weight (BM)-matched Foxj1-Cre; LRP1*_loxP/loxP_* mice at 20 weeks of age. Mice were administered IP leptin (0.5 mg/kg) and sacrificed 30 minutes later. Tissue lysates (10 µg) were separated by SDS–PAGE. Leptin and LRP1β bands were visualized by immunoblotting. The graph shows densitometric quantitation of immunoblot leptin from LRP1*_loxP/loxP_* and Foxj1-Cre; LRP1*_loxP/loxP_* (n = 3 per group). All bars and errors represent means ± SEM. *P* values by two-sided student’s *t*-test. \*\**P* < 0.01 vs. LRP1*_loxP/loxP_* mice by two-sided student’s t-test.

**Supplemental Fig. 5.**
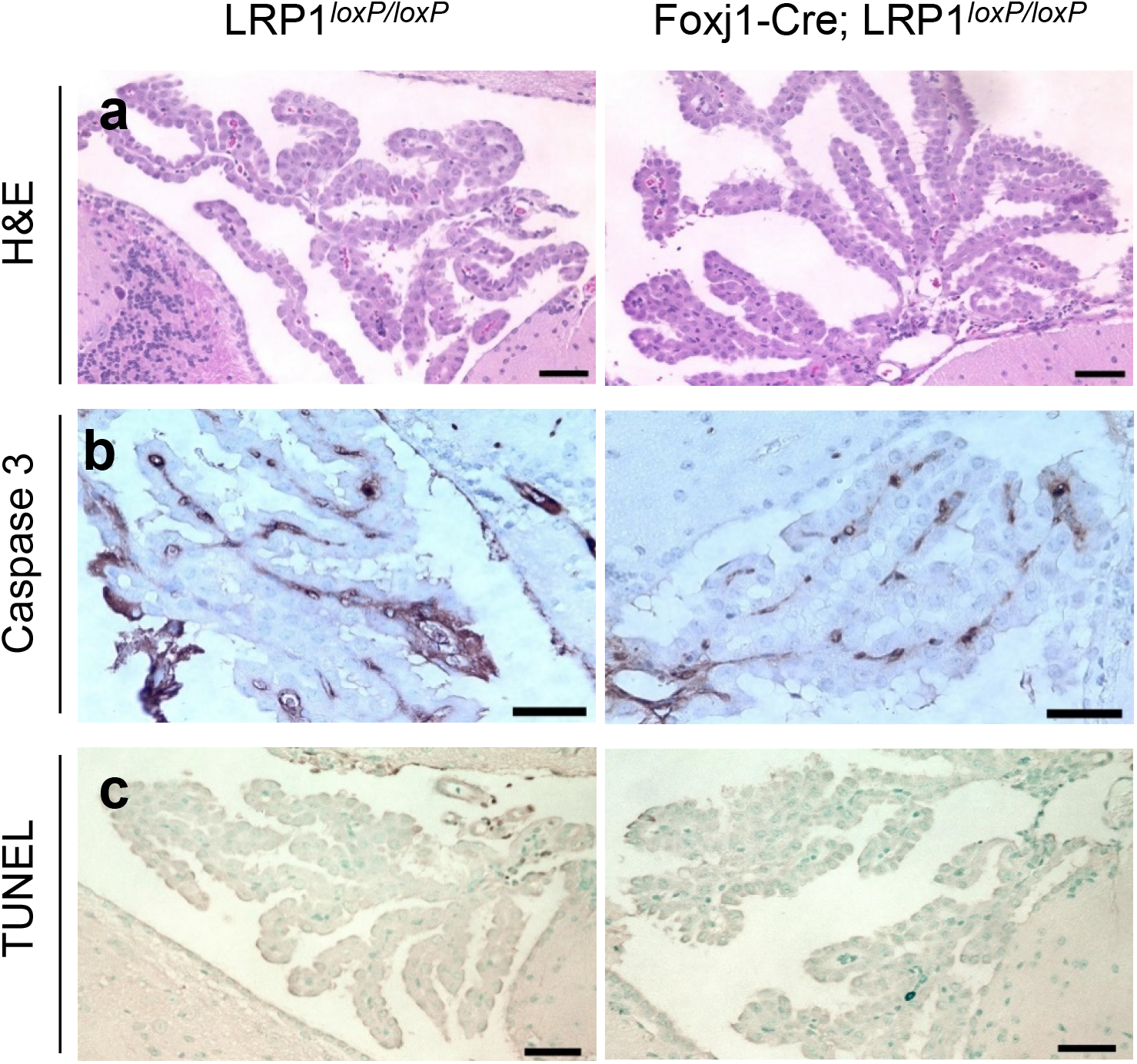
Histopathological features in the choroid plexus (ChP) from male LRP1*_loxP/loxP_* and Foxj1-Cre; LRP1*_loxP/loxP_* mice at 16 weeks of age. (a) Histologic H&E staining in the ChP between LRP1*_loxP/loxP_* and Foxj1-Cre; LRP1*_loxP/loxP_* mice. (b) Immunohistochemical analysis of caspase 3 in the ChP of LRP1*_loxP/loxP_* and Foxj1-Cre; LRP1_l*oxP/loxP*_ mice. (c) TUNEL assay in the ChP between LRP1*_loxP/loxP_* and Foxj1-Cre; LRP1*_loxP/loxP_* mice. The scale bars represent 30 µm.

**Supplemental Fig. 6.**
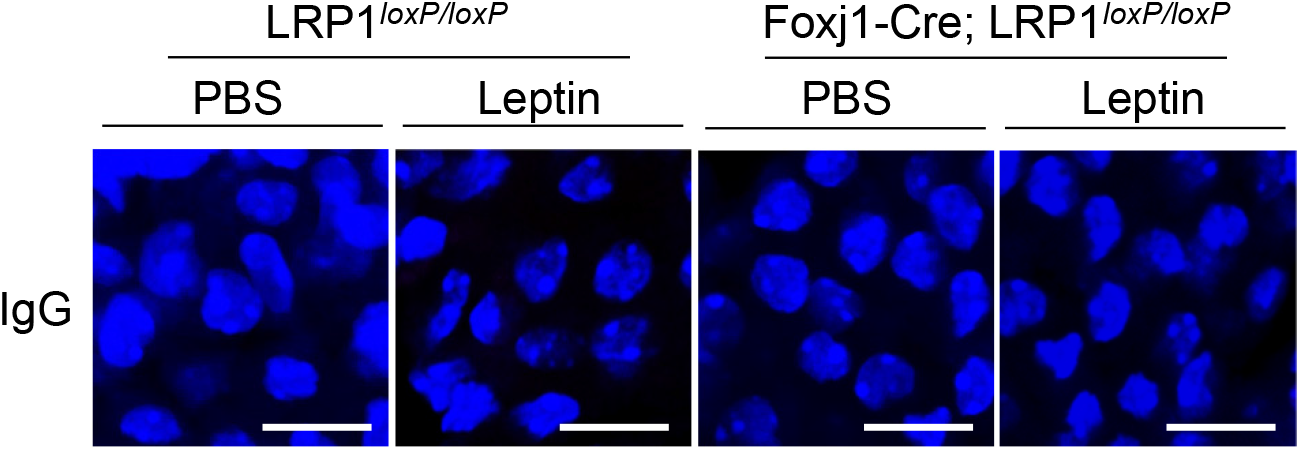
Leptin induces the physical interaction of LRP1 and LepR in epithelial cells of the choroid plexus *in vivo*. LRP1*_loxP/loxP_* and Foxj1-Cre; LRP1*_loxP/loxP_* mice at 18 weeks of age were administered with IP leptin (1 mg/kg) and sacrificed 30 min later. The choroid plexus samples were harvested, and proximity ligation assays was performed with both normal rabbit IgG and normal mouse IgG. The scale bar represents 20 μm.

**Supplemental Fig. 7.**
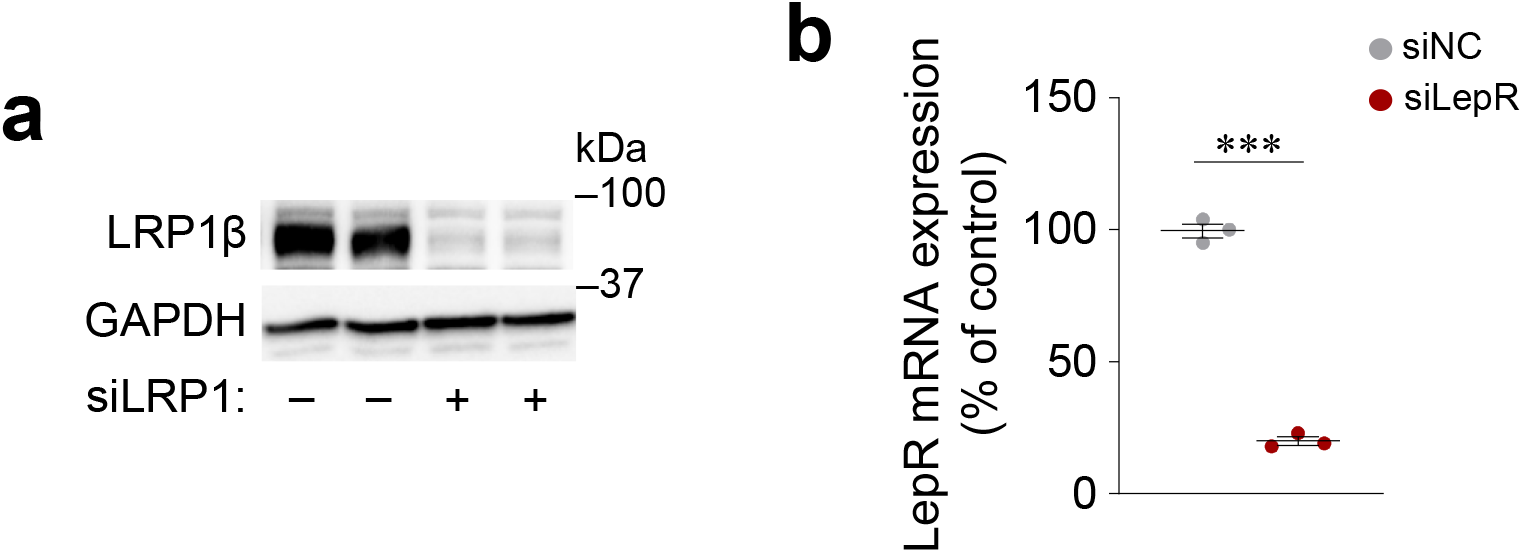
Expression of LRP1 or LepR by siRNAs in rat choroidal plexus epithelial Z310 cells (Z310 cells). (a) Rat choroidal plexus epithelial Z310 cells (Z310 cells) were transiently transfected with siRNA for negative control (siNC) or siRNA for LRP1 (siLRP1) for 48 h. LRP1β and GAPDH bands were visualized by immunoblotting. (b) Z310 cells were transiently transfected with siRNA for LepR (siLepR) or siNC for 48 h. The mRNA levels of LepR and β-Actin were measured by *q*RT-PCR. All bars and errors represent means ± SEM. \*\*\**P* < 0.001 vs. siNC by two-sided student’s *t*-test.

**Supplemental Fig. 8.**
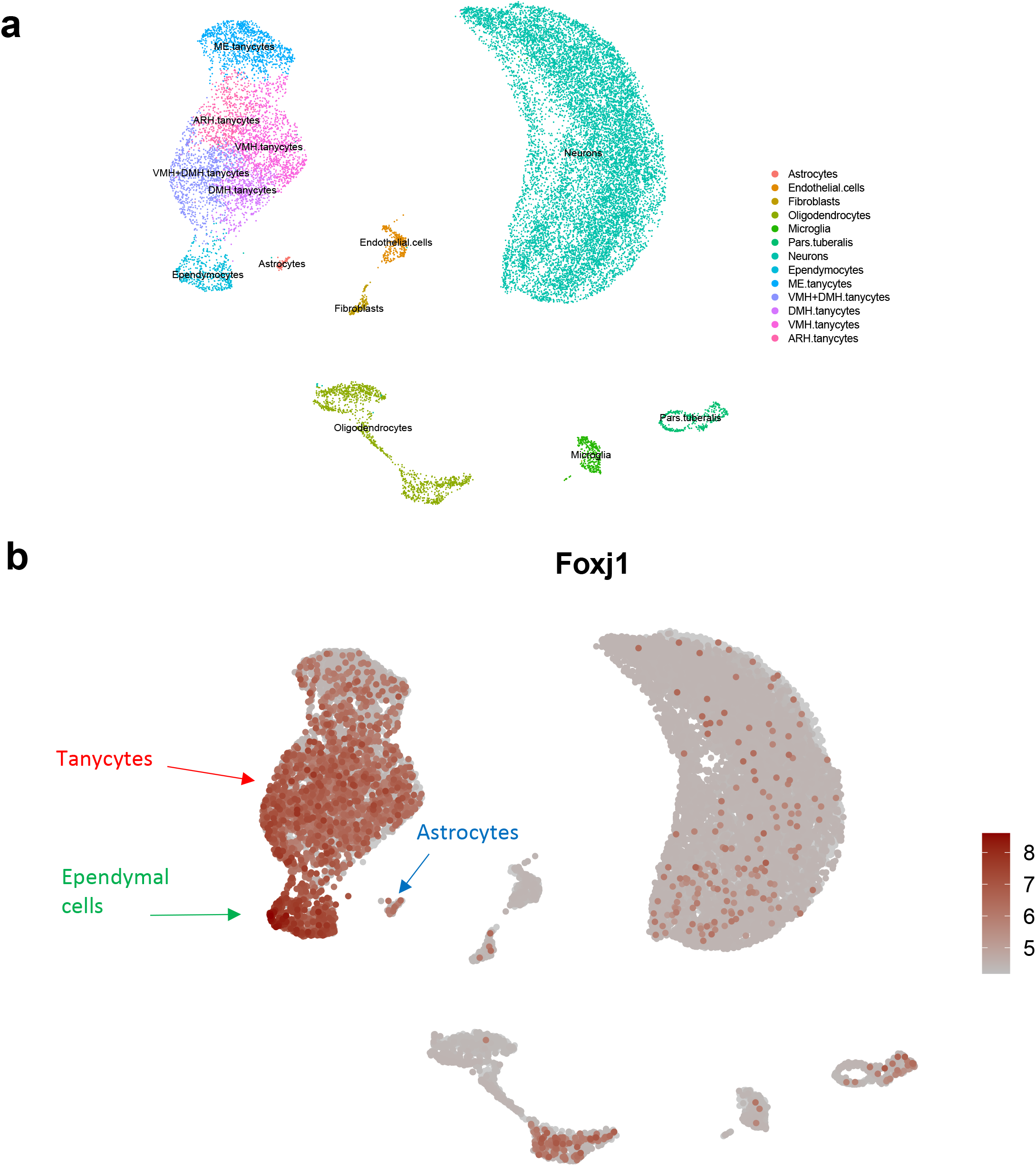
Enrichment of *Foxj1* expression in tanycytes in the ventral periventricular region of the tuberal hypothalamus (a) UMAP plot of 20,921 cells in the mouse arcuate nucleus of the hypothalamus and median eminence (PMID: 28166221). (b) Feature plot showing the expression of *Foxj1* in the arcuate nucleus of the hypothalamus and the median eminence.

